# Data-driven RNA phenotyping captures genetically regulated dimensions of the transcriptome

**DOI:** 10.64898/2026.02.20.707100

**Authors:** Daniel Munro, Alexander Gusev, Abraham A. Palmer, Pejman Mohammadi

## Abstract

Transcriptomic diversity across individuals arises from multiple modes of RNA regulation—including pre-mRNA expression, splicing, degradation, and other processes—and has been widely leveraged to map quantitative trait loci (xQTLs) and interpret GWAS signals. We recently developed a multimodal framework called Pantry that can extend discovery beyond total expression by integrating multiple transcriptomic modalities. However, Pantry and similar tools remain limited by their reliance on complete gene annotations and the statistical complexity of jointly analyzing correlated modalities. Here, we present LaDDR (Latent Data-Driven RNA phenotyping), a mechanism-agnostic framework that generates orthogonal, latent coverage features per gene, enabling xQTL discovery and GWAS integration without requiring complete gene annotations. Applied to GTEx, LaDDR identified an average of 95% more independent xQTLs per tissue than the six transcriptional regulation modes implemented in Pantry (“knowledge-driven”). Residualizing known modalities prior to LaDDR and combining with knowledge-driven phenotypes increased discovery by an additional 41% per tissue on average, while retaining the interpretability of knowledge-driven signals. In a transcriptome-wide association study (TWAS) of 114 complex traits, using LaDDR-derived phenotypes uncovered an average of 11,790 unique gene–trait pairs per tissue, versus 8,579 from knowledge-driven phenotypes. The newly captured genetic signals exhibit functional and colocalization qualities consistent with known mechanisms, suggesting that LaDDR broadens the detectable landscape of trait-relevant transcriptomic regulation by efficiently recovering regulatory variation missed by current pipelines.

## Introduction

Transcriptome sequencing data have been used to characterize variation in transcriptome regulation across individuals^1–4^. Integration with genome variation data has been used extensively to understand the regulatory mechanisms that mediate disease risk and variation in complex traits^5–8^. While these analyses were originally introduced using gene expression data, over the years several other modes of transcriptome regulation, such as splicing, the use of alternative transcript start and end sites, and others, have further expanded the scope of transcriptome analysis^9–12^. The conventional approach to transcriptome analysis involves building specific tools that are designed to quantify each specific mode of transcriptome regulation in an RNA-seq library based on the mechanistic knowledge.

While this *knowledge-driven phenotyping* approach maximizes interpretability, it has several limitations. First, it relies on potentially inaccurate annotations of transcriptome products and regulatory events specific to each mode of regulation. This complicates phenotype extraction because unannotated regulatory events are either excluded or reported inappropriately under known annotation entries^13^. Notably, a study of 90 GTEx samples for which long-read sequencing data were available identified 70,000 unannotated transcript isoforms for annotated genes in the human genome^14^. Conversely, not all annotated transcript isoforms can be accurately quantified from short read sequencing data. Another limitation of knowledge-driven phenotyping is the pervasive correlations within the data, which lead to false discoveries, redundant findings, and a loss of statistical power. These include both biological correlation, technical artifacts in quantification, and systematic correlation across different modes of transcription regulation^15^.

Finally, incorporating multiple modes of transcriptome regulation into a single analysis is often too cumbersome due to practical constraints, including the need to harmonize and run many packages and the need for domain-specific expertise. Tools such as the eQTL-Catalogue/rnaseq pipeline^16^ and Pantry^17^ aim to alleviate this issue by curating and harmonizing different transcriptome phenotyping software to extract multiple phenotypic modalities from RNA-seq data. Pantry currently quantifies 6 modalities: expression, isoform ratio, intron excision ratio, alternative TSS, alternative polyA, and RNA stability.

*Data driven phenotyping* approaches can address all three of these limitations by efficiently capturing variation in population transcriptome data without relying on detailed annotations. Other annotation-agnostic methods have been introduced previously to address this challenge for certain use cases. For example, SCISSOR uses base-level RNA-seq coverage to detect outliers in read depth within each gene^18^. Another method, derfinder, identifies base-level differential expression from RNA-seq coverage data^19,20^.

Here we introduce LaDDR (Latent Data-Driven RNA phenotyping), which extracts latent orthogonal dimensions of variation for each gene, serving as transcriptional phenotypes from RNA-seq data, thereby increasing its utility for QTL analysis and GWAS integration. We show that LaDDR’s data-driven phenotypes yield more xQTL and TWAS associations than Pantry’s six knowledge-driven modalities combined. Furthermore, we show that LaDDR can be used in conjunction with conventional phenotyping tools to provide even greater discovery power while retaining data interpretability. We provide a software package, pre-built models for 49 human tissues, and data including data-driven phenotypes and results of their application to genetic analyses.

## Results

### Overview of the method

The full LaDDR phenotyping procedure has four main steps (**Figure 1a**, **Methods**):

1. **Get RNA-seq coverage.** RNA-seq reads for a set of samples from one or more tissues are mapped to the genome and converted to base-level coverage. The products of these standard processing steps are often precomputed and freely available for download, for example through recount3^21^, even for datasets whose raw data are access-controlled.
2. **Partition genes into bins.** Each gene region is partitioned into bins used to summarize RNA-seq coverage. These bins are defined to segment the genome based on variation in RNA-seq coverage, grouping neighboring bases with similar coverage patterns across the reference data. We discuss different binning approaches in a subsequent section.
3. **Bin and normalize coverage.** RNA-seq coverage is averaged per bin per sample, and then normalized.
4. **Transform to data-driven phenotypes.** Principal component analysis (PCA) is applied to those values per gene, and the sample projections onto each principal component (PC) are saved as data-driven phenotypes (DDPs). These DDPs are comparable to knowledge-driven RNA phenotypes (KDPs), such as the multimodal set of phenotypes produced by Pantry, in that a number of phenotypes are produced per gene (**Figure 1b**). This allows many of the same applications of KDPs, such as genetic association methods like xQTL mapping and multimodal TWAS (xTWAS), which can operate on multiple features per gene (**Figure 1c**).

**Figure 1.**
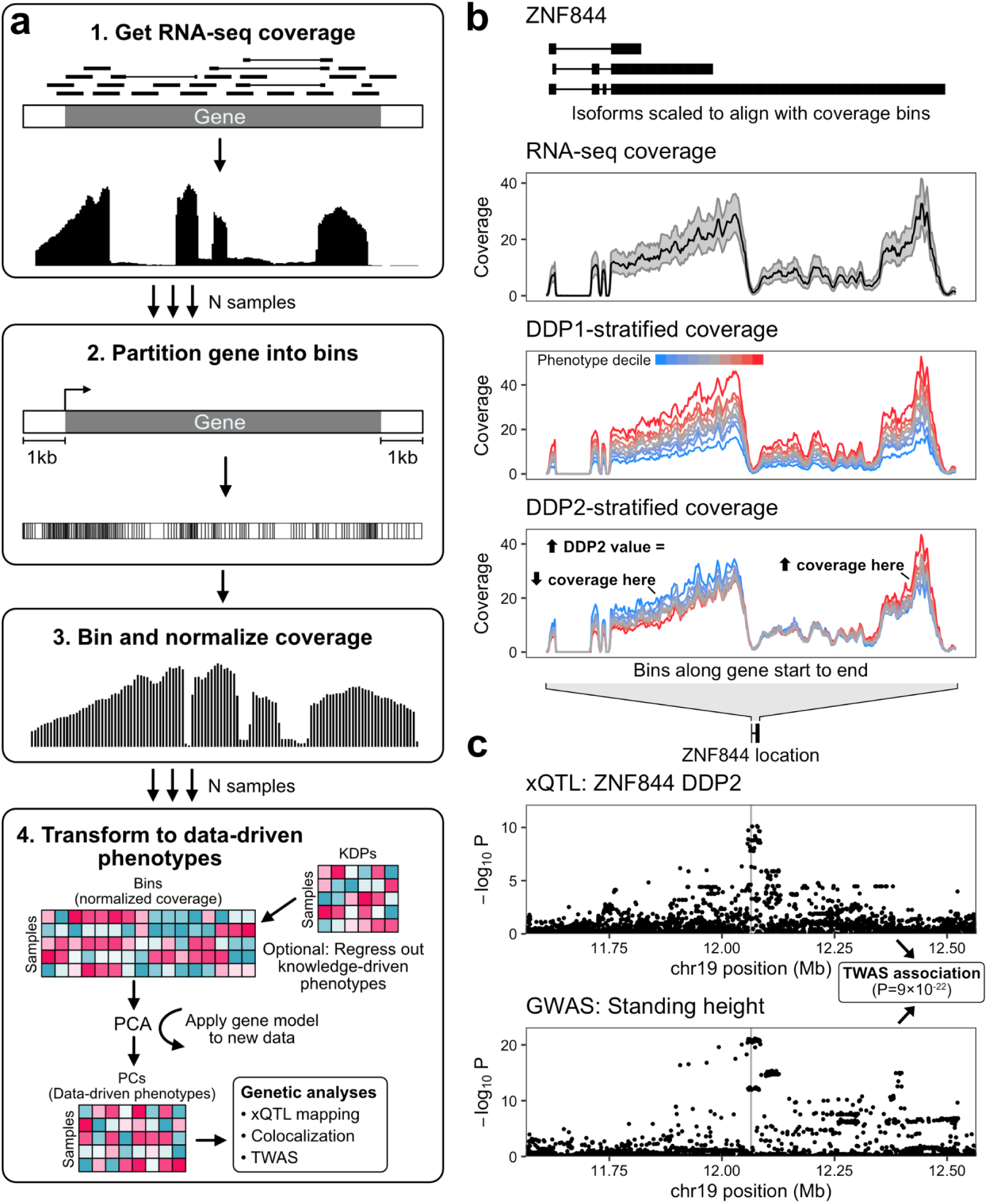
Overview of LaDDR phenotyping. **a** Diagram of the LaDDR procedure. **b** Example of DDPs for one gene, *ZNF844*. Top: The three annotated isoforms of *ZNF844* are shown, and, as in the two panels below, the x-axis represents the coverage bin index along the gene and is not proportional to genomic position, since actual bin widths vary. Isoforms are shown here for reference and are not used by LaDDR. Middle: RNA-seq coverage distribution (median and interquartile range) at each bin across all tibial nerve samples. Bottom: DDPs were generated using LaDDR. Tibial nerve samples were grouped into 10 equal-sized deciles based on their values for two resulting *ZNF884* phenotypes, DDP1 and DDP2, and median coverage at each bin is shown for each decile. Coverage in all three panels was scaled per sample to adjust for sequencing depth. **c** TWAS association for DDP2 in b. Top: Nominal *p*-values for each variant in the +/- 500 kb cis-window from xQTL mapping of *ZNF844* DDP2 in tibial nerve are shown. The location of *ZNF844* is shown above for reference and its transcription start site is indicated in the plots with a vertical line. Bottom: GWAS *p*-values for standing height (UK Biobank) in the same window.

The PCA models can then be applied to processed RNA-seq coverage data of additional datasets, and the resulting DDPs quantified for each gene will correspond (i.e., represent the same dimensions of transcriptomic variation) across all datasets generated from the same set of models. The coverage bin definitions can also be reused to generate new models on new data.

### Benchmarking strategy and optimization of LaDDR parameters

We implemented multiple strategies for segmenting gene regions into bins to summarize coverage data. These include fixed number or width of equal-sized bins per coding and noncoding region of each gene, as well as data-driven methods that aim to divide regions with greater coverage variation into more bins (**Supplementary Methods**). We ran each of these binning methods using a variety of density parameters and mapped and quantified independent cis-xQTLs on the resulting DDPs to compare their effect (see **Supplementary Methods**). In general, the data-driven binning methods based on coverage variance performed best in terms of producing the most xQTLs, and making efficient use of fewer bins (**Supplementary Figure 1a**).

In the top-performing approach, which has been set as the default and used for the results presented in this study, the RNA-seq coverage values for a gene are log_2_-transformed, the difference in coverage between consecutive bases is computed, and the variance of these differences is computed at each base. Bin boundaries are then determined such that the cumulative sum of variance differences within each bin reaches a predefined threshold. This produces a higher density of smaller bins in gene regions and genes with more transcriptional variation than other regions and genes.

By default, principal component analysis (PCA) is applied to the binned normalized coverage data per gene, and the sample projections onto each principal component (PC) are saved as data-driven phenotypes (DDPs). Functional PCA (FPCA) is also implemented as an alternative to PCA (**Supplementary Methods**). We tried different forms of FPCA, as well as PCA with different maximum numbers of PCs. The PCA runs produced more xQTLs than the FPCA runs, and as the maximum PC threshold increased, the number of resulting xQTLs increased and then decreased (**Supplementary Figure 1b**). We therefore chose PCA with up to 16 PCs per gene as the default.

### LaDDR phenotypes capture familiar and novel variation

We trained LaDDR gene models using 28,637 samples from 54 human tissues from the Genotype-Tissue Expression Project^22^ (GTEx) and 33 human cancer types from The Cancer Genome Atlas^23^ (TCGA). These datasets provide RNA-seq coverage patterns across a diverse set of healthy and dysregulated tissue contexts. We applied the models to generate DDPs for each of 49 GTEx tissue datasets, producing an average of 417,593 (SD 9,274) phenotypes per tissue across an average of 40,147 (SD 1,351) protein-coding and lncRNA genes. We also trained and applied models on 445 human lymphoblastoid cell line samples from the Geuvadis dataset^2^.

We examined the DDPs in terms of their correlation with knowledge-driven phenotypes (KDPs) generated by Pantry. While some DDPs correlate primarily with a single KDP, such as total expression, more commonly they show varying degrees of correlation with multiple KDPs for the same gene (**Figure 2a**). We also assessed cis-heritability and found that it is, on average, highest for DDP1, the first principal component for each gene, and decreases with increasing PC rank (**Figure 2b**). This pattern has a practical implication, as it supports reducing dimensionality by retaining only the leading DDPs. The cis-heritability of the earlier (lower-numbered) DDPs are comparable in distribution to the KDPs, while later (higher-numbered) DDPs have lower cis-heritability (**Figure 2c**).

**Figure 2.**
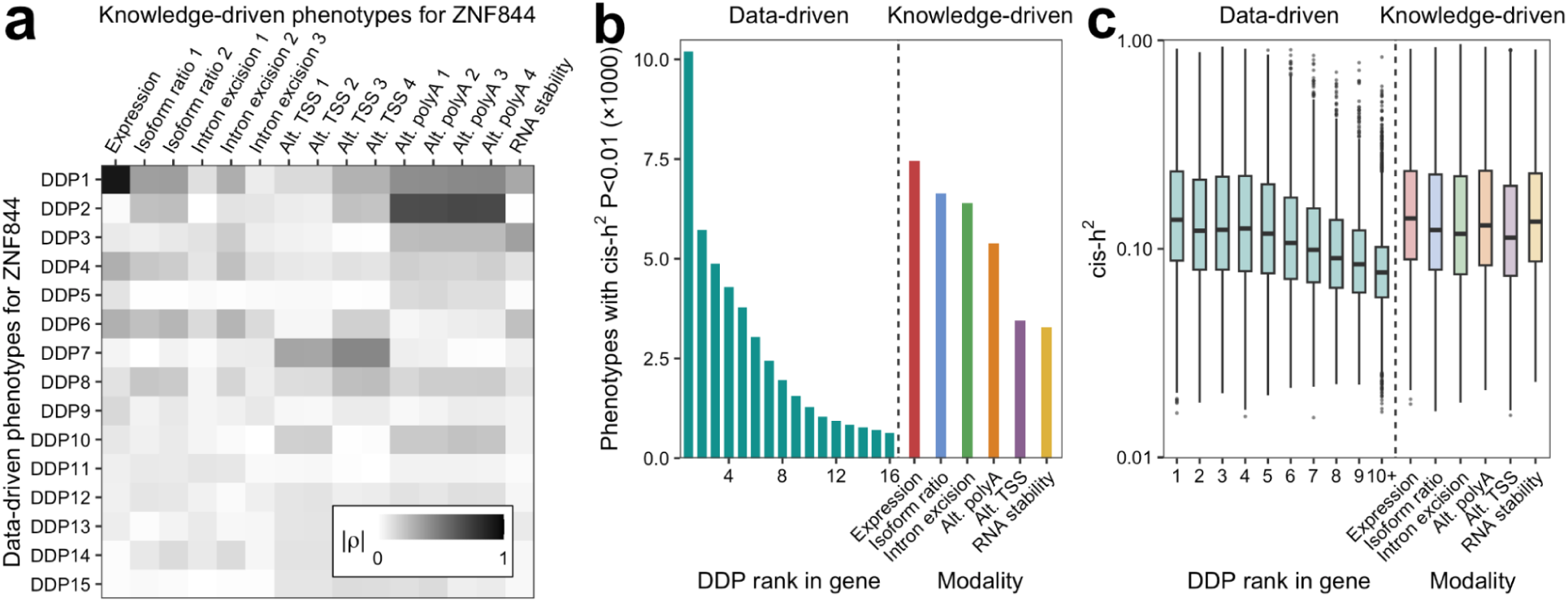
LaDDR phenotypes are similarly heritable to knowledge-driven phenotypes. **a** Heatmap of absolute value of Spearman correlations between each DDP (rows) and each KDP (columns) for one gene, *ZNF844*, across the Geuvadis dataset. **b** Number of DDPs per DDP rank and KDPs per modality in the Geuvadis dataset that were used for xTWAS mapping. Only phenotypes with heritability *p*-value <0.01 are used for xTWAS. **c** Boxplots show cis-heritability of those heritable DDPs and KDPs.

### LaDDR phenotypes encode multimodal genetic regulatory variation

We mapped cis-xQTLs (hereafter referred to as xQTLs) for the DDPs in each GTEx tissue using tensorQTL^24^ in grouped phenotype mode. We found 3,755 to 64,106 conditionally independent xQTLs per tissue for 3,419 to 29,007 genes (**Supplementary Table 1**). We compared these results with those from Pantry, in which KDPs from six modalities were jointly mapped using cross-modality mapping, finding that, on average, 95% more independent cis-QTLs were identified for DDPs than for KDPs per tissue (**Figure 3a, 3b**).

**Figure 3.**
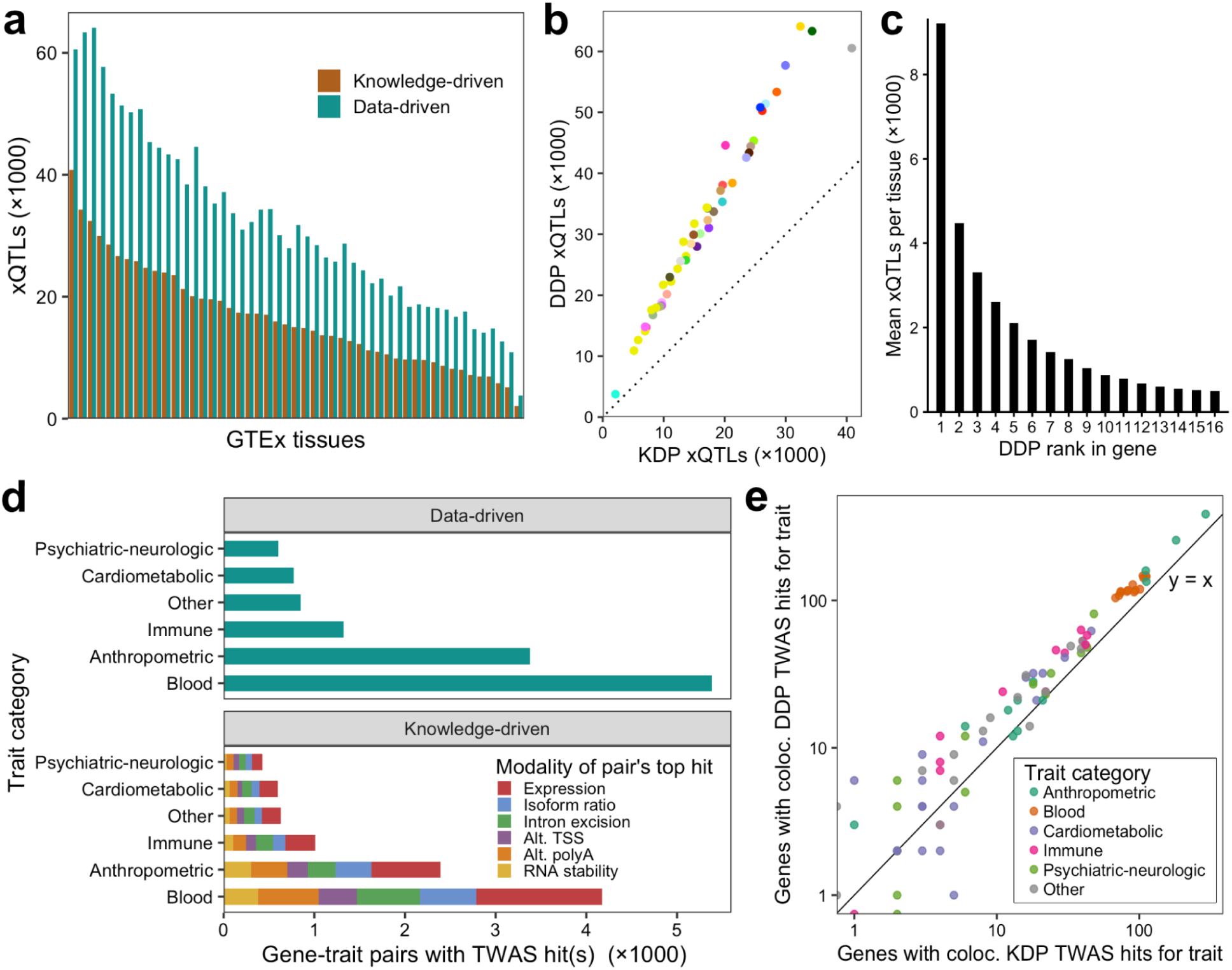
LaDDR increases cis-xQTL and xTWAS discovery. **a** xQTLs for DDPs and for KDPs in each GTEx tissue. **b** The same values shown as a scatter plot, using the tissue colors provided by GTEx. **c** Number of xQTLs, averaged over tissues, mapped for each within-gene DDP rank. **d** Unique gene-trait pairs with xTWAS hits per trait category using DDPs (top) and KDPs (bottom). **e** Number of unique gene-trait pairs with at least one colocalizing TWAS hit using DDPs (y-axis) versus KDPs (x-axis) for the transcriptomic models. Points are shown for the 86 tested traits with at least one colocalizing hit, and points with zero colocalizing hits for one of the axes are shown along an axis line to include them on the log-scaled axes.

DDPs corresponding to lower principal components of each gene PCA model tend to map more xQTLs than those corresponding to higher PCs, with DDP1 producing 7.4 times as many xQTLs as DDP8 and 18.6 times as many xQTLs as DDP16 (**Figure 3c**).

We applied DDPs as transcriptomic models for xTWAS. Similar to the multimodal approach previously reported using Pantry phenotypes^17^, we used FUSION to fit a transcriptomic prediction model to each DDP in each tissue and then applied them to GWAS data for a collection of 114 complex traits^7^. Compared to six-modality xTWAS at the same *p*-value threshold of 5⨉10^-8^ and using the same Geuvadis dataset for transcriptomic models, DDPs resulted in nearly the same number of total associations (24,697 vs. 24,644 for six-modality), 33% more unique gene-trait pairs with associations (**Figure 3d**), and 37% more unique gene-trait pairs with strong evidence of colocalization at the level of shared causal variants (**Figure 3e**). Applying xTWAS to DDPs for 49 GTEx tissues and the same 114 traits resulted in a median of 26,501 significant TWAS associations per GTEx tissue, including a total of 64,314 unique gene-trait pairs, 28,116 of which include strongly colocalizing associations.

### LaDDR phenotypes can complement knowledge-driven phenotypes for genetic discovery

We developed a variant of the LaDDR phenotyping method that produces residual data-driven phenotypes (rDDPs), designed to complement KDPs and serve as a “catch-all” modality alongside them. This is accomplished by regressing out all KDPs from the normalized coverage features prior to PCA on a per-gene basis (see **Methods**). We produced rDDPs for Geuvadis using Pantry’s six default modalities of KDPs and examined the maximum Pearson r^2^ between each rDDP and a KDP of the same gene. For PC, residualization reduced these values from mean 0.40 (SD 0.27) to mean 0.068 (SD 0.093) (**Figure 4a**). For all PCs, residualization reduced the max r^2^ values from mean 0.072 (SD 0.14) to mean 0.030 (SD 0.055).

**Figure 4.**
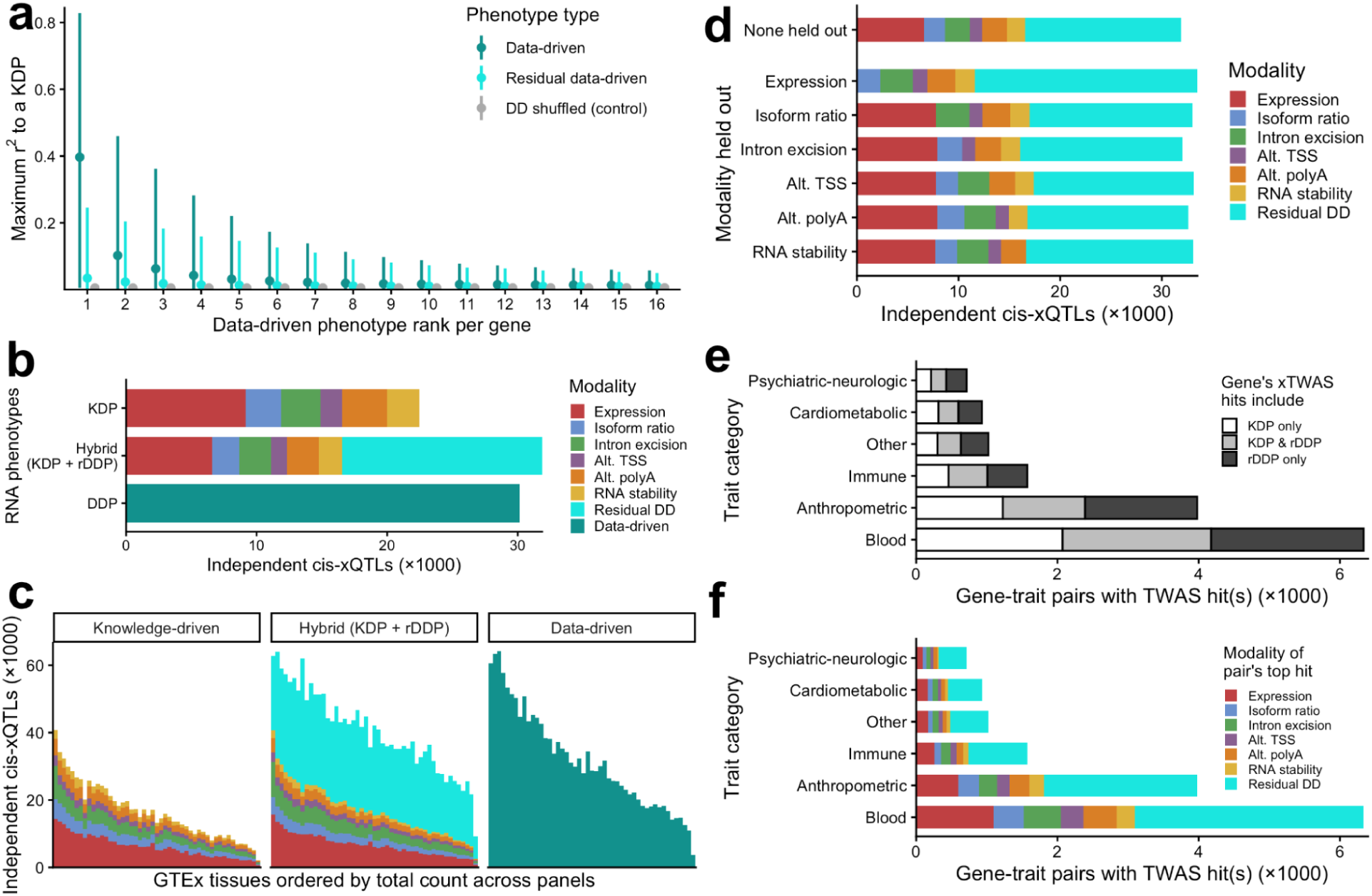
Residual data-driven RNA phenotypes complement knowledge-driven phenotypes. **a** Correlations between data driven (full and residual) and knowledge-driven RNA phenotypes for Geuvadis. As a control, values for each DDP were shuffled and maximum r^2^ with KDPs were recomputed. **b** Independent xQTLs produced by cross-modality mapping on six modalities of KDPs, cross-modality mapping on six modalities of KDPs plus rDDPs, and cis-QTL mapping on only DDPs for Geuvadis. **c** Independent xQTLs from the same three forms of cross-modality mapping applied to each GTEx tissue. **d** xQTLs from hybrid (KDP + rDDP) cross-modality mapping using all six KDP modalities, and xQTLs resulting from hybrid cross-modality mapping where each modality of KDPs was individually held out from rDDP generation and xQTL mapping. **e** Number of unique gene-trait pairs with hybrid (KDP + rDDP) xTWAS hits per trait category. Pairs are colored by whether the associated phenotypes per gene-trait pair included one or more KDPs, one or more rDDPs, or both. **f** The same gene-trait pairs as e, but grouped and colored by the modality of the hit with the lowest *p*-value.

To compare the ability of different sets of RNA phenotypes to represent cis-regulatory variation, we mapped cis-xQTLs that are conditionally independent across all phenotypes per gene using Pantry’s cross-modality mapping procedure. We applied this to KDPs, DDPs, and a hybrid set of phenotypes (KDPs + rDDPs) for the Geuvadis dataset. Mapping KDPs resulted in 22,470 total xQTLs, while mapping DDPs resulted in 30,158 xQTLs. Adding rDDPs to the set of KDPs resulted in a net increase of 9,408 xQTLs (42%) for a total of 31,878, 16,544 of which were called for KDPs and 15,334 of which were called for rDDPs (**Figure 4b**). In terms of unique genes represented by the xQTLs, the addition of the rDDPs increased the number of xGenes from 12,872 to 16,950. Repeating the KDP, DDP, and hybrid xQTL mapping procedures on each GTEx tissue produced consistent results in terms of the relative number of independent xQTLs (**Figure 4c**, **Supplementary Table 1**).

To assess the ability of the rDDPs to capture modalities of transcriptomic variation that are not explicitly measured, we repeated the residual phenotyping procedure, each time withholding one KDP modality and residualizing normalized coverage data on the other five. We then repeated hybrid cross-modality mapping, each time using phenotypes for only five KDP modalities plus the rDDPs residualized on those five modalities. In every case, holding out a single modality increased the number of rDDPs in compensation, yielding a similar total number of independent xQTLs (**Figure 4d**).

We examined the additional rDDP xQTLs obtained by holding out each KDP modality. Since the exact correspondence of xQTLs across different xQTL mapping results is not straightforward, we examined only rDDP xQTLs for genes that lacked rDDP xQTLs in the non-held-out hybrid mapping. As expected, the positions of these xQTLs relative to their xGene somewhat reflect the modality that was held out (**Supplementary Figure 2**).

We then examined the application of rDDPs to xTWAS. Unlike cross-modality mapped xQTLs, conditional independence across phenotypes and modalities is not enforced by the TWAS method used in this study, so we ran TWAS on the rDDPs for Geuvadis and simply combined the results with those from KDPs. This resulted in a 58% increase in unique gene-trait association pairs (**Figure 4e**), and rDDPs showed the strongest associations for 7,655 (53%) of the pairs (**Figure 4f**). Similarly, across GTEx tissues, adding rDDP TWAS hits increased the average number of unique gene-trait association pairs by 59% (**Supplementary Table 2**), and rDDPs had the strongest associations for 6,926 (51%) of the pairs on average.

Of particular interest are gene-trait pairs that have xTWAS hits for both DDPs and rDDPs, but no hits for KDPs in any tissue. Four such gene-trait pairs are highlighted here. The “Morning/evening person (chronotype)” trait had data-driven xTWAS hits for *CCNH* (Cyclin H) in adrenal gland tissue. *CCNH* was previously mapped to a GWAS locus for chronotype^25^. The “Triglycerides” trait had data-driven xTWAS hits for *RFLNA* (Refilin A) in left ventricle tissue.

*RFLNA* was previously mapped to a GWAS locus for triglyceride levels^26^. The “Fasting glucose” trait had a data-driven xTWAS hit for *DGKB* (Diacylglycerol kinase beta) in pancreas. *DGKB* was previously mapped to a GWAS locus for type 2 diabetes mellitus^27^. Finally, the “Sleeplessness / insomnia” trait had a data-driven xTWAS hit for *LINC03051* (Long intergenic non-protein coding RNA 3051) in the anterior cingulate cortex (BA24). *LINC03051* was previously mapped to a GWAS locus for insomnia^28^. These examples corroborate prior GWAS discoveries and demonstrate how gene-trait genetic associations missed by knowledge-driven phenotyping can be captured by LaDDR phenotyping.

### Functional characteristics of phenotypes and their genetic associations

We aggregated the positions of KDP, rDDP, and DDP xQTLs relative to their xGenes and found that the distributions of both rDDP and DDP xQTLs exhibited a mixture of the major features of the KDP xQTLs, namely concentration within the gene body with peaks at the gene start and end (**Figure 5a**). DDP xQTLs, which were mapped on their own, showed a stronger peak at the gene start, while rDDP xQTLs, which were residualized using KDPs and included in hybrid cross-modality mapping alongside KDPs, had a lower peak at the gene start than at the gene end.

**Figure 5.**
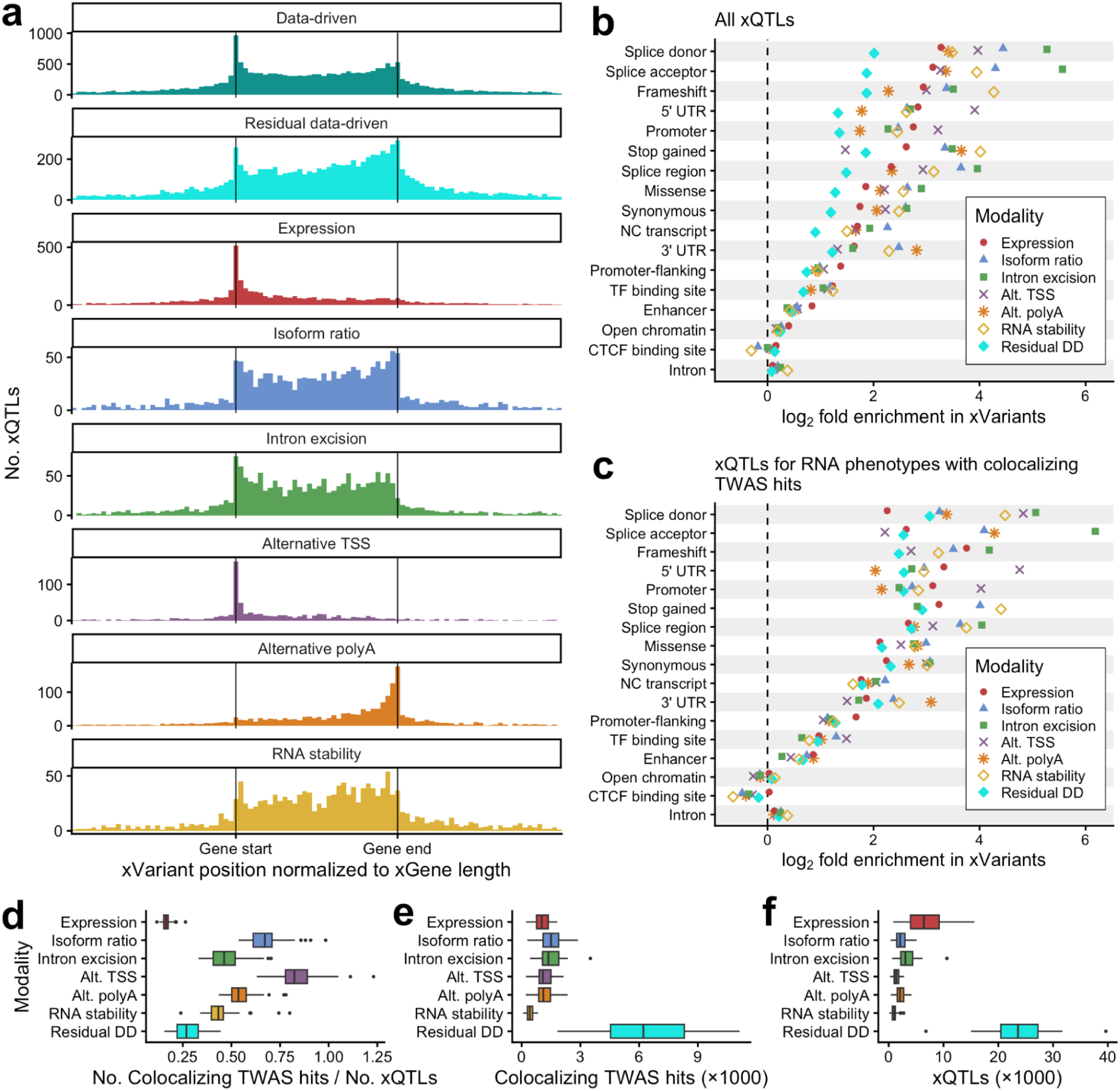
Functional characteristics of LaDDR phenotyping. **a** Histograms of DDP, rDDP, and KDP xQTL locations relative to their xGene. The first panel shows xQTLs from DDP xQTL mapping, and the remaining seven panels show xQTLs from hybrid xQTL mapping across all KDPs and rDDPs. The genomic coordinates of each xGene and that gene’s xQTLs were linearly transformed such that the gene starts and ends are aligned on the x-axis. xQTLs more than one gene length from the gene are not shown. Histograms are colored by modality. **b** Enrichment of functional categories in the fine-mapped xQTLs from hybrid cross-modality mapping. For each modality of xQTLs, log_2_-ratio of the prevalence of each annotation among fine-mapped xQTLs to the background prevalence is shown. Enrichment was computed for xQTLs aggregated across 49 GTEx tissues. Categories are sorted by expression modality enrichment value. **c** Enrichment was computed the same way as in b, but after filtering xQTLs to those for RNA phenotypes with at least one colocalizing TWAS hit in the same tissue. **d** For each modality, the number of colocalizing xTWAS hits, divided by the number of xQTLs from hybrid cross-modality mapping, was computed for each GTEx tissue and shown as a boxplot. **e** The numerators for the ratios plotted in d are shown as boxplots. **f** The denominators for the ratios plotted in d are shown as boxplots.

We then measured enrichment of various functional annotations in the fine-mapped xQTLs of each modality. For most functional categories, rDDP xQTLs were enriched, but less so than any knowledge-driven modality (**Figure 5b**). However, when filtering xQTLs to those for RNA phenotypes that also had colocalizing xTWAS hits in the same tissue, functional enrichment in rDDP xQTLs was similar to that of knowledge-driven modalities (**Figure 5c**).

We also compared the relative rate of colocalization with GWAS loci per xQTL for rDDPs and KDPs. Expression had the lowest ratio of colocalizing TWAS hits to xQTLs (median 0.16 across tissues), and rDDPs had the second-lowest (median 0.27, **Figure 5d-f**). Expression and rDDPs also had the lowest ratios for unique genes with colocalizing TWAS hits per unique xQTL gene and for colocalizing TWAS hits per xQTL when filtered to the top hit per gene-trait pair and top xQTL per gene (**Supplementary Figure 3**).

To check for indications that DDPs could be more susceptible to false positives due to misassignment of RNA-seq coverage to nearby genes, we measured the rate at which genes in close proximity were associated with the same trait, noting that such cases can also have other causes, such as pleiotropy and genuine independent mediators. Defining nearby as having any overlap according to the gene’s transcription start and end positions in the gene annotations, 35.6% of DDP TWAS gene-trait pairs for protein-coding genes had a nearby gene associated with the same trait, compared to 31.4% of KDP TWAS pairs (**Supplementary Figure 4**). For gene-trait pairs involving lncRNAs, the percentages were higher, at 59.9% for DDP TWAS and 57.7% for KDP TWAS. The similarity of these percentages for DDPs versus KDPs suggests that misassigned RNA-seq coverage is not substantially higher in LaDDR phenotyping than in the more established RNA phenotyping methods.

Despite long non-coding RNAs (lncRNAs) having higher transcript complexity (i.e., splice variants per exon) than protein-coding mRNAs^29^, xQTL mapping across six KDP modalities produced an average of 0.16 independent xQTLs per gene per tissue for lncRNAs, which is only 29% of the rate for protein-coding genes. Multiple biological and methodological factors could be contributing to this difference, but gene annotation completeness could be one factor. In the annotations used in this study, lncRNAs have 28.9% the number of exons per gene summed over isoforms compared to protein-coding genes. However, DDP xQTL mapping showed less of a difference, with 0.39 xQTLs per lncRNA per tissue, which was 44% of the rate for protein-coding genes.

### Impact of input data quality

Data driven phenotypes from LaDDR are robust across datasets, library types and are minimally impacted by missing annotations. We quantified this robustness by running LaDDR using different amounts of training data, simulated sparse reference annotations, and simulated variation in input RNA-seq read types and lengths.

Because some transcriptomic variation is tissue-specific, we hypothesized that fitting LaDDR gene models using more samples from more tissues and other contexts would result in phenotypes that capture more regulatory variation. We fitted a set of models using five GTEx tissues from different tissue categories (Adipose - Subcutaneous, Artery - Aorta, Brain - Frontal Cortex (BA9), Esophagus - Gastroesophageal Junction, and Skin - Sun Exposed (Lower leg)), a separate set of models using all 54 GTEx tissues, and a third set of models using all 54 GTEx tissues and 33 cancer types from The Cancer Genome Atlas (TCGA)^23^. We compared them by using each set to generate DDPs for 49 GTEx tissues and map cis-xQTLs for each tissue for each set of models. Unexpectedly, using 54 rather than five tissues to fit models increased the number of independent xQTLs per tissue by only 4.2% on average (**Supplementary Figure 5a**). Using 54 tissues plus 33 cancer types increased xQTLs by 1.7% on average compared to 54 tissues only (**Supplementary Figure 5b**). In other words, xQTLs for a tissue can be found at nearly the same rate regardless of whether that particular tissue was used to fit the LaDDR models.

To measure the effect of gene annotation sparsity on the ability for DDPs to represent regulatory variation, we generated a series of increasingly sparse annotations by dropping non-canonical isoforms in random order until only the canonical isoform for each gene remained. We then used each set of gene annotations to generate DDPs using LaDDR and KDPs using Pantry, and then mapped xQTLs for the phenotypes. DDPs were less affected by gene annotation sparsity than were KDPs, with independent xQTLs dropping 1.8% between 100% and 0% inclusion of non-canonical isoforms, compared to a 14% drop for KDPs (**Supplementary Figure 6**).

To estimate the effect of RNA sequencing technologies on LaDDR phenotyping quality, we selected 100 Geuvadis samples, which have 75 bp paired-end RNA-seq, and altered the reads in three ways. These included truncating all reads to 50 bp, treating paired-end reads as single-end, and subsampling reads to fractions of the original depths. When we used each of these variations as input for LaDDR and mapped cis-QTLs to the DDPs, we observed a relatively small drop (-2.7%) in discoveries from the 50 bp truncated reads, and larger drops from the single-end (-7.2%) and lowered sequencing depth (-41% for 50% of reads, -66% for 25% of reads) simulations (**Supplementary Figure 7**).

## Discussion

We introduced LaDDR, a method for data-driven RNA phenotyping that efficiently captures regulatory variation with minimal dependence on reference gene annotations. We showed that DDPs encode more regulatory variation than six major modalities of knowledge-driven RNA phenotypes combined, both in terms of the number of independent cis-xQTLs and the number of gene-trait xTWAS associations discovered. We also described residual data-driven phenotypes that, when combined with a set of knowledge-driven phenotypes, increased the discovery of independent cis-xQTLs by 42% and the number of gene-trait pairs with xTWAS associations by 37%. While data-driven phenotypes are generally less functionally interpretable than KDPs, DDPs trade off this limitation with the advantage of encoding more cumulative genetic regulatory variation, while rDDPs serve the purpose of being used in combination with KDPs to maintain interpretable genetic associations with KDPs and supplement them with additional discoveries.

Though LaDDR joins existing methods such as SCISSOR^18^ and derfinder^20^ in extracting features from base-level RNA-seq coverage data, it was built to enable direct application to population genetic analyses. Thus, rather than focusing on outliers or differential expression, it produces molecular phenotypes that can be genetically mapped in the same way as knowledge-driven molecular phenotypes.

While some methods for latent embeddings of genomic data use deep learning^30–32^, the approach introduced here uses features that are linearly related to log-transformed expression coverage. One advantage of this approach is a straightforward characterization of phenotypes by examining the PCA loadings for the coverage bins across the gene, revealing which portions of the gene vary in proportion to that phenotype’s values. Another advantage is that the PCs represent orthogonal dimensions of variation, providing a compact set of features that minimize redundancy in their resulting genetic associations and overfitting.

Two of the presented analyses together suggest why LaDDR discovers more gene regulatory associations than the panel of KDPs used in this study. The analysis in which KDPs were held out of rDDP residualization and xQTL mapping one modality at a time (**Figure 4d**) showed that DDPs from LaDDR can encode the regulatory variation in KDPs, so long as they are not intentionally regressed out. The analysis comparing the number of LaDDR DDP xQTLs versus KDP xQTLs using subsampled isoform annotations (**Supplementary Figure 6**) showed that LaDDR is minimally impacted by annotation sparsity. Assuming the currently known gene annotations are incomplete, these observations suggest that LaDDR DDPs encode regulatory variation that could be identified in KDPs using future annotations. However, the leveling off of KDP xQTLs in the subsampled isoform annotations analysis, far below the number of DDP xQTLs, suggests that DDPs capture regulatory variation beyond what the KDP modalities implemented by Pantry can capture, even with fully comprehensive annotations. These two interpretations suggest that DDPs can capture two forms of transcriptomic variation missing from a collection of KDPs: uncharacterized modalities of variation, and specific instances of variation in known mechanisms that were not represented in the reference gene annotations. Both of these benefits may decrease as mechanistic knowledge and transcriptome annotation increase. Still, the compactness and computational efficiency of data-driven phenotypes may remain advantageous for some applications.

We also showed that, despite relying minimally on gene annotations and no modality-specific knowledge or methodology, the use of DDPs for xTWAS resulted in more discovered gene-trait associations than xTWAS with six transcriptomic modalities using Pantry. Another advantage of using PCA-derived, data-driven phenotypes for xTWAS is the reduction of redundant associations arising from correlated phenotypes for the same gene. Still, when maximal discovery of gene-trait associations and potentially implicated modalities is a priority, a hybrid set of phenotypes (KDPs + rDDPs) can be used.

We found that the LaDDR method presented here is more effective at quantifying regulatory variation in lncRNAs than the six KDP modalities we tested. Given the high amount of genetic variation within lncRNAs and their high diversity of functions^33^, the ability to capture lncRNA regulatory variation is an important component of transcriptome-wide genetic analyses.

Nevertheless, the inclusion of non-coding transcripts poses challenges for avoiding misassignment of RNA-seq reads to nearby or overlapping gene regions. In the current LaDDR implementation, we address this by excluding regions from one gene that overlap with annotated exonic regions of another gene for the purposes of RNA-seq coverage quantification. Refinements of this procedure, such as a probabilistic method of gene assignment, could further reduce potential misassignment of coverage variation while maximizing the inclusion of coverage data for each gene. Still, our analysis of nearby genes sharing DDP xTWAS associations showed only a slight increase in such cases compared to KDP xTWAS.

For most variant functional categories, we observed less enrichment in rDDP xQTLs than in KDP xQTLs (**Figure 5b**), though this difference was diminished after filtering to RNA phenotypes with colocalizing TWAS hits (**Figure 5c**). One hypothesis is that variant annotations correspond somewhat to known gene annotations, and so RNA phenotypes residualized to remove variation quantified using known annotations will associate more with variants that are un-annotated. However, it could also indicate that rDDPs produce a higher fraction of xQTLs that are spurious in some way than do KDPs, though further research would be needed to assess these two hypotheses. A related observation was that while rDDPs exhibit a relatively low ratio of colocalizing TWAS hits to xQTLs, expression QTLs have an even lower ratio. Given that rDDPs and expression produced more xQTLs than the other modalities, their power may result in detection of a higher fraction of weaker regulatory effects that are less likely to associate strongly with complex traits.

We found that LaDDR models built with 54 tissue datasets produced slightly more xQTLs than those built with 5 tissue datasets, and adding 33 cancer datasets provided a slight additional improvement. We speculate that models trained on multiple tissues do not capture much of the tissue-specific variation patterns, especially since only genes expressed in most of the full set of training samples are included, thus omitting genes with highly tissue-specific expression. Future versions of LaDDR could explore whether fitting tissue-specific or other context-specific models results in an increase in tissue-specific, and thus an increase in total, genetic associations, and whether that benefit outweighs potential drawbacks such as lower sample size per model set, greater computational burden, and the loss of correspondence of phenotype representations across tissues.

LaDDR greatly expands the information extracted from RNA-seq data that can be applied to genetic analyses. We provide DDPs, rDDPs, and the resulting xQTLs and TWAS weights and associations in the Pantry Portal (https://pantry.pejlab.org/) alongside their knowledge-driven counterparts. We also provide the LaDDR code as a Python package (https://github.com/PejLab/LaDDR) and have integrated it into the Pantry framework (https://github.com/PejLab/Pantry) to facilitate the joint use of knowledge-driven and data-driven RNA phenotypes to enhance genetic discovery.

## Methods

### LaDDR phenotyping

The input data are base-level RNA-seq coverage files in bigWig format. Protein-coding and lncRNA genes are first partitioned into bins using an adaptive binning process. RNA-seq coverage at every base across the gene, from start to end plus 1kb upstream and downstream, is extracted for 256 randomly chosen samples from the training set. Values are log_2_-transformed after adding a pseudocount (eight by default), coverage difference at consecutive bases is computed, and variance of the differences is computed at each base. Starting at the beginning of the region, bases are included in the current bin until the cumulative sum of variance differences reaches a predefined threshold, after which the next bin begins. This produces a higher density of smaller bins in regions of the gene with more transcriptional variation. The threshold, which is fixed across all genes, is determined using a random sample of 128 genes and calculated to produce a desired average number of bins per gene (256 by default). Bins larger than 1024 bases are subdivided uniformly such that the maximum bin size is 1024. Finally, any bin overlapping an exonic region of a different gene is removed to reduce contamination of a gene’s RNA-seq coverage by reads from a different gene’s transcript.

Using the bin definitions, base-level coverage counts are summarized into mean coverage per bin per sample using pyBigWig^34^. A scaling factor is computed for each sample based on cumulative coverage level relative to the median sample level, and coverage values are divided by the scaling factors to adjust for sequencing depths. A pseudocount (eight by default) is added to the coverage values, which are then log_2_-transformed to stabilize variance.

At this stage, knowledge-driven phenotypes, such as those produced by Pantry, can be regressed out of the normalized binned coverage features to represent residual transcriptomic variation. This happens on a per-gene basis, with the following procedure applied to each bin feature using all KDPs for the same gene. First, an ordinary least squares linear regression is fit using all of the gene’s KDPs as features and normalized coverage for the bin as the response variable. KDPs with regression *p*-value <0.01 are kept, and linear regression is repeated with only those KDPs as variables. The regression model predictions on the same data are subtracted from the bin feature and the residuals are used in place of the original coverage features in subsequent steps.

The normalized binned coverage data or the residualized data are then standardized and used to fit a PCA model for each gene using the scikit-learn Python package^35^. Genes are filtered such that models are only produced for those with nonzero total coverage in at least 50% of samples. While the number of PCs is necessarily capped at the minimum of the number of bins and the number of samples, additional caps can be set to explain up to a desired percent of variance and a maximum number of PCs. By default and for this study, enough PCs to explain 80% of variance, up to a maximum of 16, were used. Genes are processed in batches for efficiency, and data can be concatenated across multiple datasets to generate models representing diverse biological contexts.

These PCA models are then applied to the same normalized binned coverage data and/or new data, one dataset at a time, and the resulting PC projections are the DDPs or rDDPs. Genes are again filtered for nonzero total coverage in at least 50% of samples, this time in the set of samples for which phenotypes are being generated.

### RNA-seq datasets

We downloaded all bigWig files for the 54 tissues from the GTEx Project (18,948 samples) and 33 cancer types from TCGA (11,287 samples) from recount3^21^. We subsetted the GTEx samples to those 17,350 included in the v8 release to have at most one sample per individual per tissue. We also used RNA-seq data for 445 human lymphoblastoid cell line samples from Geuvadis^2^. To get bigWig coverage files for Geuvadis, we mapped RNA-seq reads to the GRCh38 reference genome using STAR v2.7.11b^36^ using the parameters specified in the Pantry pipeline and then converted each BAM file to bigWig files using bamCoverage from deeptools^37^ with --binSize 1.

### Pantry data

Pantry was used to quantify knowledge-driven RNA phenotypes for comparison with data-driven RNA phenotypes and to generate residual data-driven RNA phenotypes. We ran Pantry on Geuvadis and 49 GTEx tissues using Ensembl v113 gene annotations^38^, producing KDPs for six modalities: expression, isoform ratio, intron excision ratio, alternative TSS, alternative polyA, and RNA stability. For some analyses, such as the pruned annotation comparison, Pantry was rerun, e.g., with different gene annotations, on particular tissue datasets, as described in the Methods subsections.

### Application of LaDDR phenotyping

We fitted and applied LaDDR DDP models to the combined set of 54 GTEx tissues and 33 TCGA cancer types using default parameters described above, and unless otherwise specified, these phenotypes were used for all reported analyses of GTEx DDPs. Likewise, we fit and applied LaDDR rDDP models to the 49 GTEx tissues for which we had Pantry phenotypes. DDP and rDDP models were also fitted and applied to the Geuvadis dataset. For comparisons of input data size and diversity, we fit DDP models to the combined set of 54 GTEx tissues, as well as a subset of five GTEx tissues (Adipose - Subcutaneous, Artery - Aorta, Brain - Frontal Cortex (BA9), Esophagus - Gastroesophageal Junction, and Skin - Sun Exposed (Lower leg)), chosen randomly from each of the five largest tissue site categories in terms of unique tissues.

### xQTL mapping

We performed cis-QTL mapping on DDPs using tensorQTL^24^, following the default Pantry Pheast pipeline. For the hybrid phenotype set xQTLs, rDDPs were added to the set of KDPs as a seventh modality, and cross-modality mapping was performed accordingly. For DDP xQTLs, DDPs were mapped as a single modality, i.e., mapping of conditionally independent cis-QTLs on phenotypes grouped by gene.

### TWAS

We generated xTWAS predictive models for DDPs and rDDPs and tested xTWAS associations using FUSION^6^ with default parameters, as described previously^17^. We tested associations for a collection of 114 GWAS traits^7^ [https://zenodo.org/record/3629742#.XjCh9OF7m90]. We used a genome-wide *p*-value threshold of 5×10^-8^ to determine significant xTWAS hits. We also performed xTWAS for KDPs generated by Pantry using the same procedure and *p*-value threshold of 5×10^-8^ to compare to the DDP xTWAS results and to combine with the rDDP xTWAS results. Colocalization tests as implemented in the coloc R package^39^ were applied to each TWAS hit, and hits with COLOC.PP4 (posterior probability of shared causal variant) > 0.8 were considered colocalizing TWAS hits for the purposes of this study.

### Pruned annotation analysis

Starting with the Ensembl human v113 gene annotations, the isoform per gene that was present (ignoring Ensembl ID version) in a curated list of canonical isoforms from UCSC [http://hgdownload.soe.ucsc.edu/goldenPath/hg38/database/knownCanonical.txt.gz] were labeled as canonical. To simulate sparser gene annotations, non-canonical isoforms were subsampled successively to keep 80%, 60%, 40%, 20%, and 0% of the original amount, with 0% leaving only the one canonical isoform per gene. Each GTF file was then used to generate DDPs using LaDDR and rDDPs using Pantry for one GTEx tissue, Brain frontal cortex (BA9) (N=175 genotyped samples). We mapped conditionally independent cis-QTLs for each set of phenotypes using tensorQTL and compared the numbers of cis-QTLs.

### Read length and depth simulations

We used a random subsample of 100 Geuvadis samples for all read length and depth simulations. We used seqtk trimfq [https://github.com/lh3/seqtk] to truncate all (75 bp) reads in the paired-end FASTQ files to 50 bp. We used seqtk sample to subsample reads in each FASTQ file to 50% and 25% of the original number, using a fixed random seed so that read pairs were preserved within R1/R2 file pairs. We then aligned reads in each simulated dataset using STAR, generated bigWig coverage files using bamCoverage, ran LaDDR, and mapped cis-QTLs on the DDPs using tensorQTL, all in the same way as described above.

### Method and parameter comparisons

To compare the performance of different variations of the LaDDR procedure, we generated DDPs for the Geuvadis dataset using each variation of methods and parameters (**Supplementary Methods**), for only protein-coding genes on chr7 for efficiency. We then mapped cis-QTLs for each of these phenotype sets using tensorQTL and used the number of conditionally independent cis-QTLs as a performance metric.

### Functional enrichment analysis of fine-mapped xQTLs

For all RNA phenotypes with xQTLs from hybrid cross-modality mapping in each of 49 GTEx tissues, we applied SuSiE^40^ fine-mapping via tensorQTL in cis_susie mode on the same input data. We measured functional enrichment on variants in the 95% credible sets for xQTLs of each modality, combined across all tissues. We downloaded variant annotations from the GTEx Portal [https://storage.googleapis.com/adult-gtex/references/v8/reference-tables/WGS_Feature_overlap_collapsed_VEP_short_4torus.MAF01.txt.gz]. For the credible set variants of each modality, the fraction of variants in the credible sets annotated with each category was computed, weighted by posterior inclusion probability (PIP), and divided by the fraction of background variants annotated. The background set included all variants in the +/- 1Mb cis-windows of phenotyped genes. Enrichment values for which the weighted count was <0.1 were excluded due to low information.

## Data availability

The LaDDR phenotypes, xQTLs, and xTWAS results for GTEx and Geuvadis data presented in this study are available at https://www.dropbox.com/scl/fo/57wjrmwqs9wwn3pd9n3xr/ANXEvS8VlPUkzCb3P2zNcmE?rlkey=bpqaww1cf4722iri0pkfkqawu&st=r1aaf6ce&dl=0 [will be replaced with Zenodo link upon acceptance]. Transcriptomic model weights for running xTWAS are available for all DDPs, rDDPs, and KDPs for GTEx tissues and Geuvadis at https://pantry.pejlab.org/download. GWAS summary statistics used for xTWAS are available at Zenodo record 3629742 [zenodo.3629742]^41^.

## Code availability

LaDDR is implemented as a Python package available at https://github.com/PejLab/LaDDR.

Code used to run LaDDR and other analyses for this study is at https://github.com/daniel-munro/laddr-cluster. Code for final data processing and generation of figures, tables, and statistics is at https://github.com/daniel-munro/laddr-paper.

## Acknowledgements

Funding: National Institute on Drug Abuse [P30DA060810, A.A.P. and U01DA060443, P.M.].

