## Supplementary Information for "Data-driven RNA phenotyping captures genetically regulated dimensions of the transcriptome"

**In this file:**
Supplementary Methods
Supplementary Tables 1-2 (descriptions)
Supplementary Figures 1-7

### Supplementary Methods

#### Comparison of binning methods and parameters

We compared the default binning method to four alternative methods, all currently available in the LaDDR software. Each method has one or more parameters that generally control the number and size of bins, so we tested the following ranges of parameter values for each method:

- **Fixed number of bins per gene subregion**:
  - Take the union of all exon regions for any isoform of a gene to get contiguous exonic regions, and split each region into a fixed number of equal-sized bins. Also split each region between adjacent exonic regions, as well as the 1kb upstream and downstream regions, into the same number of bins.
  - Range of parameter values compared: 4-32 bins per region.
- **Fixed bin widths**:
  - Using the same regions as above, split each coding (exonic) region into bins of a fixed width, and split each non-coding region into bins of a different fixed width chosen to be eight times larger than coding region bins.
  - Range of parameter values compared: 16/128 to 64/512 for coding/non-coding bin widths.
- **Data-driven binning based on cumulative coverage and coverage correlation**:
  - Start with each base as its own bin and merge bins in two stages:
    1. While any bin has mean cumulative coverage per sample below a threshold, merge it with whichever of its two neighbors has lower cumulative coverage, and repeat for the next lowest-coverage bin.
    2. Measure correlation between neighboring bins as Pearson correlation of $\log_2$(norm. coverage + 1) across samples. Normalized coverage means, for each sample, the fraction of cumulative coverage across the gene that is within the bin. While any neighboring bins have correlation above the threshold, merge them and repeat.
  - Range of parameter values compared: 16-128 minimum cumulative mean coverage per bin, 0.8-0.95 maximum correlation between adjacent bins.
- **Data-driven binning based on cumulative variance of coverage**:

    - Determine the cumulative variance threshold: Get variance of $\log_2$(coverage + 8) at each base. Get sum of variance across each gene (for a sample of 128 genes for efficiency), get mean sum per gene, and divide by desired number of bins per gene to get sum of variance per bin. Spikes of variance can skew the resulting number of bins per gene away from the intended target, so the following procedure is performed once on the sample of 128 genes and the threshold is adjusted to accurately produce the desired number of bins per gene. Binning is then performed for all genes.
    - For each gene, start with the first base and extend the bin until the cumulative-variance-per-bin threshold is reached. Repeat until reaching the end of the gene's range.
    - Range of parameter values compared: 128-1024 mean bins per gene.
- **Data-driven binning based on cumulative variance of coverage diffs (the default)**:
    - Identical to the previous method, but compute the differences in adjacent values for $\log_2$(coverage + 8) at each base, and calculate variance of those (rather than on $\log_2$(coverage + 8) directly) to determine the cumulative variance threshold and the bin boundaries.
    - Range of parameter values compared: 128-1024 mean bins per gene.

For each set of parameter values for each method, we ran the full phenotyping procedure with otherwise default parameters (e.g., PCA models with a maximum of 16 PCs per gene) on the Geuvadis dataset. We then performed cis-QTL mapping on the DDPs to use independent cis-QTLs as a metric for each binning's ability to capture cis-regulatory variation. For efficiency, we computed DDPs and cis-QTLs only for chr7 (932 genes).

#### Comparison of model types and parameters

We compared the use of PCA to functional PCA as the LaDDR phenotyping model. While basic PCA uses the normalized binned coverage values as an unordered set of input variables, functional PCA can use the relative position of the bins along the gene sequence as additional information. We compared the following variants of PCA and FPCA:

- **PCA**: For each gene, keep enough PCs to capture at least 80% variance explained, then clipped to a fixed maximum number of PCs. Range of parameter values compared: 6-24 maximum PCs kept.
- **Discretized FPCA**: Use the `scikit-fda` Python package, providing the binned coverage features in order.
- **FPCA with spline basis using bin coordinates**: Use a 4th-order BSpline basis, using ordered bin numbers as domain coordinates.
- **FPCA with spline basis using equal bin spacing**: Use a 4th-order BSpline basis, using the middle coordinate of each bin as domain coordinates.

As with the binning comparison, we generated DDPs for the Geuvadis dataset using each of these variants with otherwise default parameters (e.g., data-driven binning based on cumulative variance of coverage diffs with average bins per gene of 256), and then mapped cis-QTLs for chr7 genes.

### Supplementary Tables

**Supplementary Table 1**: xQTL count per tissue for DDP, hybrid (KDP + rDDP), and KDP phenotype sets

**Supplementary Table 2**: xTWAS hit and gene counts per trait per tissue for DDP, rDDP, and KDP phenotype sets

### Supplementary Figures

**Supplementary Figure 1: Comparison of parameters and methodological variations.**

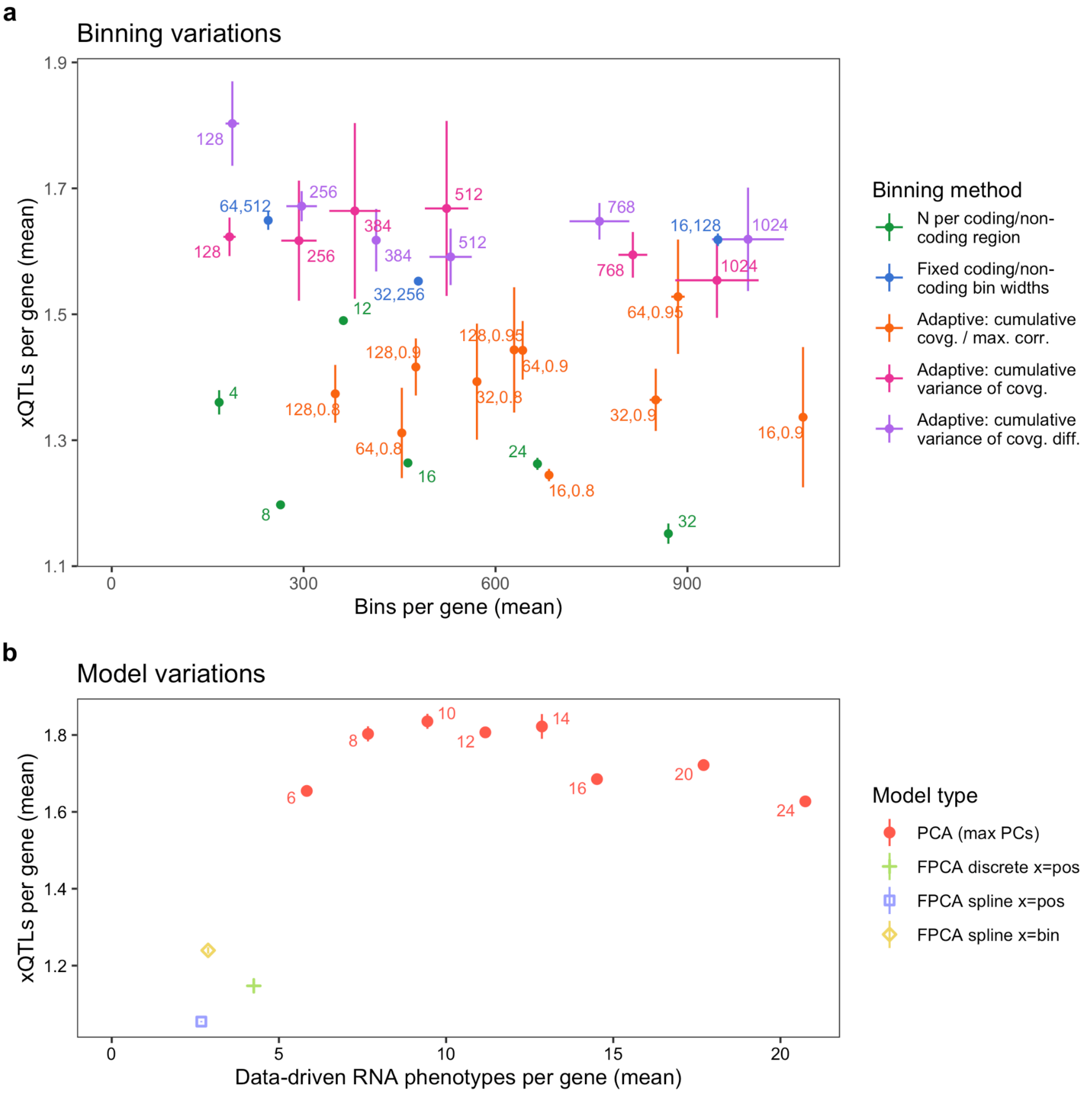


**a** Average number of bins per gene and conditionally independent DDP cis-xQTLs per gene for each binning method variation. Points and lines represent mean and standard deviation, respectively, across three trials for each method-parameter pair. Points are labeled with method-specific parameters as follows, following the order in the legend: number of bins per coding/noncoding region, bin widths for bins inside coding and noncoding regions, minimum cumulative mean coverage per bin and maximum correlation between adjacent bins, mean number of bins per gene, and mean number of bins per gene. **b** Average number of DDPs per gene and conditionally independent cis-xQTLs per gene resulting from each LaDDR gene model variation. Points for PCA models are labeled with the maximum number of PCs kept per gene.

**Supplementary Figure 2: New rDDP xQTLs when holding out specific knowledge-driven modalities.**

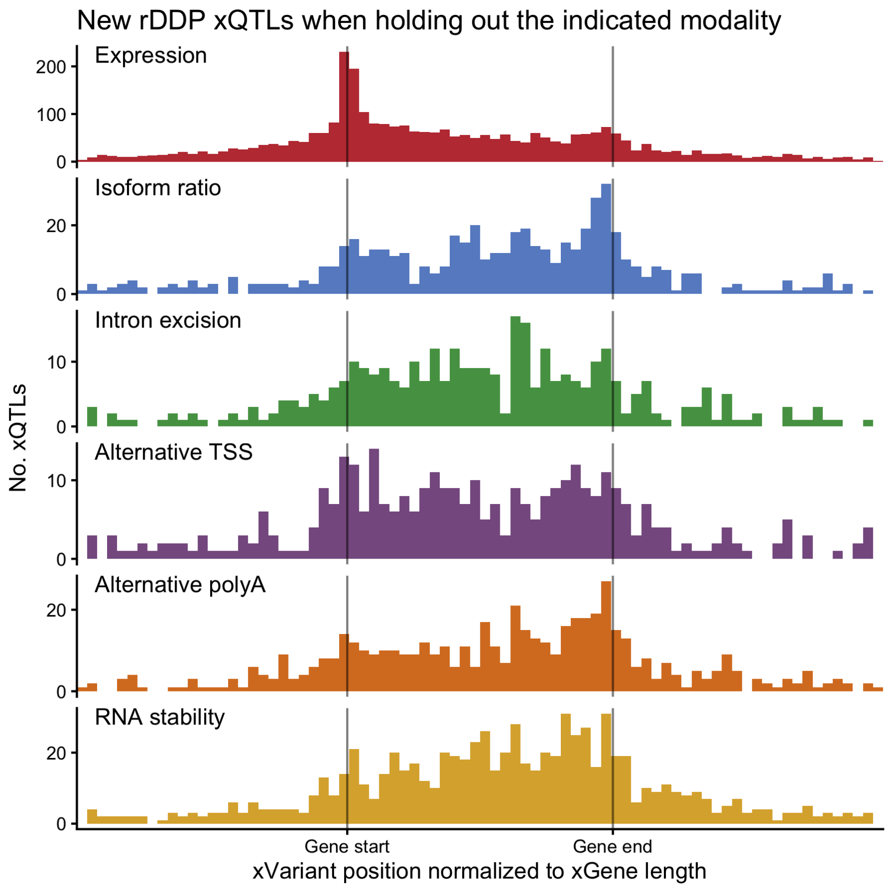


Histograms of rDDP xQTL locations relative to their xGene. The process of generating rDDPs and doing cross-modality mapping with KDPs and rDDPs was repeated six times, each time withholding one modality from the residualizing procedure and the xQTL mapping. Each xQTL shown is for an rDDP generated by holding out the indicated KDP modality, excluding those in genes that also had an rDDP xQTL in the non-held-out run. The genomic coordinates of each xGene and that gene's xQTLs were linearly transformed such that the gene starts and ends are aligned on the x-axis. xQTLs more than one gene length from the gene are not shown. Histograms are colored by modality.

#### Supplementary Figure 3: Additional colocalization to xQTL ratios

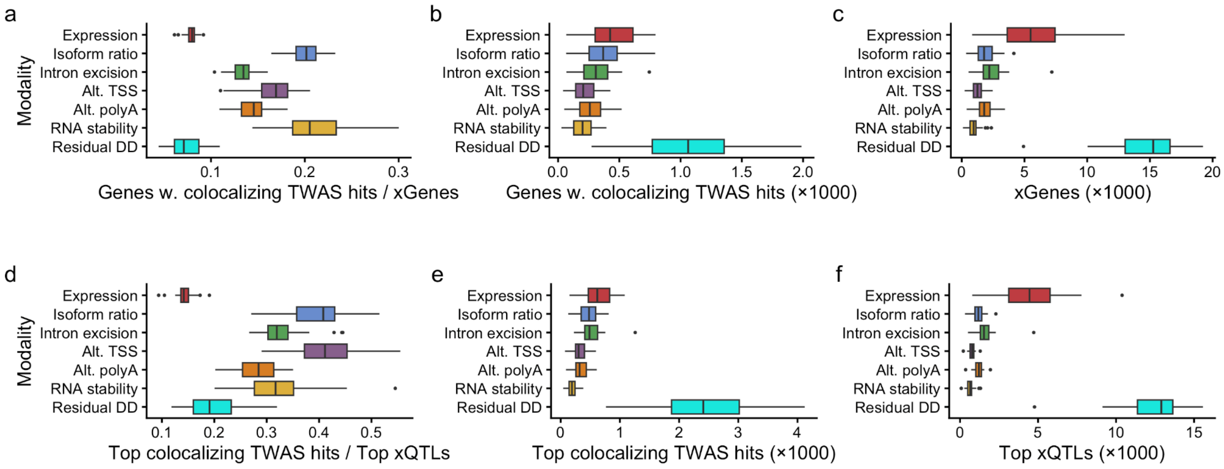


**a** For each modality, the number of genes with one or more colocalizing (COLOC.PP4 >0.8) xTWAS hits, divided by the number of genes with one or more xQTLs from hybrid cross-modality mapping, was computed for each GTEx tissue and shown as a boxplot. **b** The numerators for the ratios plotted in a are shown as boxplots. **c** The denominators for the ratios plotted in a are shown as boxplots. **d** For each tissue, the strongest colocalizing TWAS hit in terms of TWAS *p*-value per gene-trait pair was selected, and the number of those belonging to each modality was counted. Likewise, for each tissue, the strongest xQTL per gene from hybrid cross-modality mapping was selected, and the number of those belonging to each modality was counted. The ratio between these values are plotted as boxplots. **e** The numerators for the ratios plotted in d are shown as boxplots. **f** The denominators for the ratios plotted in d are shown as boxplots.

**Supplementary Figure 4: Percentage of xTWAS hits shared with nearby genes**

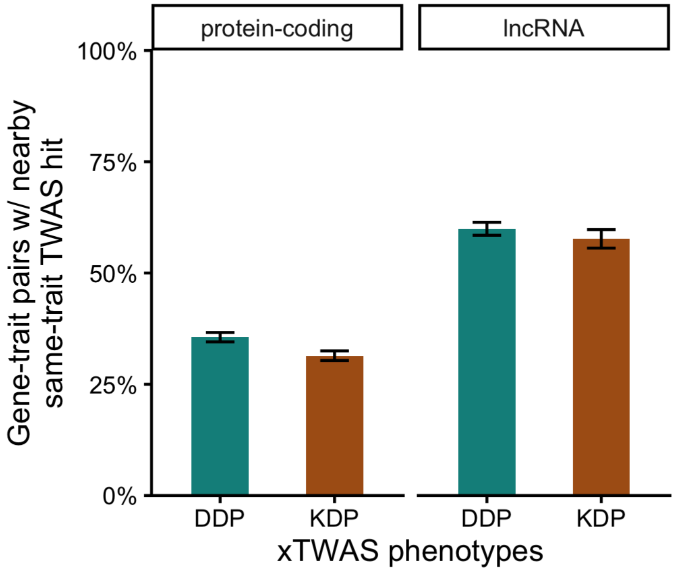


For each unique gene-trait pair with at least one DDP xTWAS hit, we examined whether there is a DDP xTWAS hit for the same trait for any nearby gene, defined as a gene whose range between transcription start and end overlaps with the first gene's range. We repeated this for KDP xTWAS hits and in both cases calculated the means separately for protein-coding and lncRNA genes. Error bars show 95% confidence intervals of each proportion. DDP = knowledge-driven phenotype, KDP = data-driven phenotype.

**Supplementary Figure 5: Performance of LaDDR models using different training data**

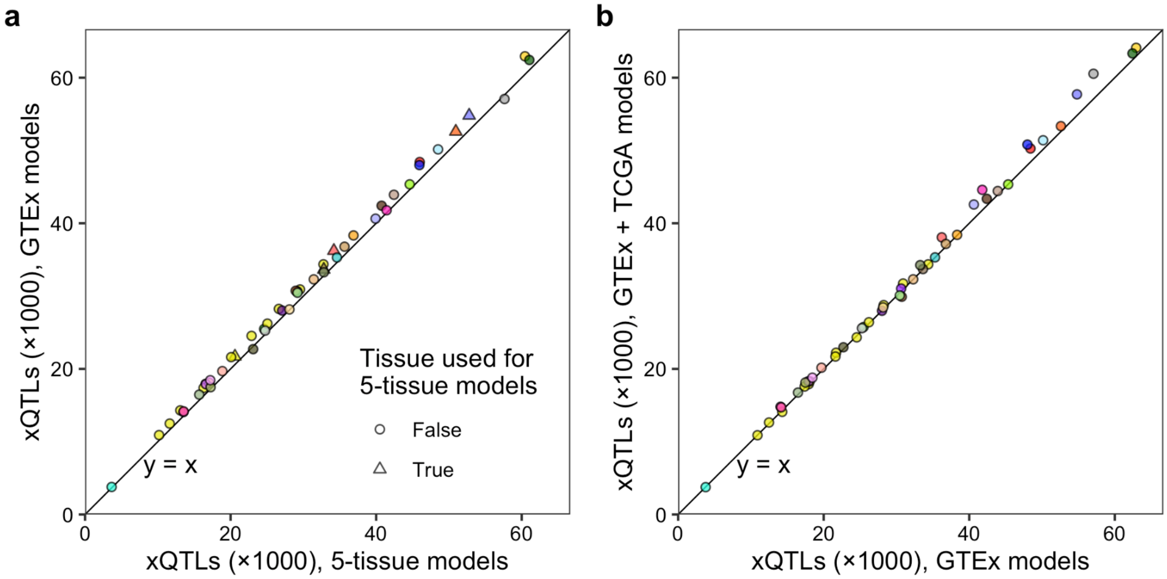


**a** Number of DDP xQTLs produced for each of 49 GTEx tissues when either all 54 GTEx tissues or only five GTEx tissues are used to fit the LaDDR gene models. **b** Number of DDP xQTLs produced for each of 49 GTEx tissues when either all 54 GTEx tissues plus 33 cancer types or only the 54 GTEx tissues are used to fit the LaDDR gene models.

**Supplementary Figure 6: xQTLs from pruned annotation simulation**

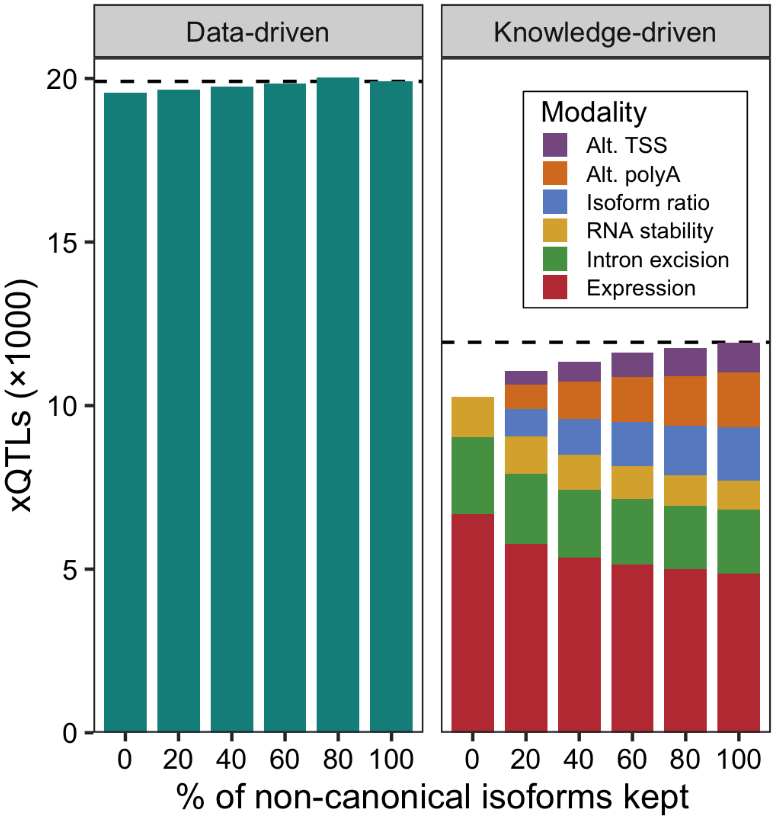


Number of xQTLs from DDPs (left) or KDPs (right) for the Brain - Frontal Cortex (BA9) tissue (N=175) after randomly subsampling non-canonical isoforms from the gene annotations used for phenotyping. Dashed lines mark the counts for 100% of isoforms for reference.

**Supplementary Figure 7: Simulated sequencing data variations**

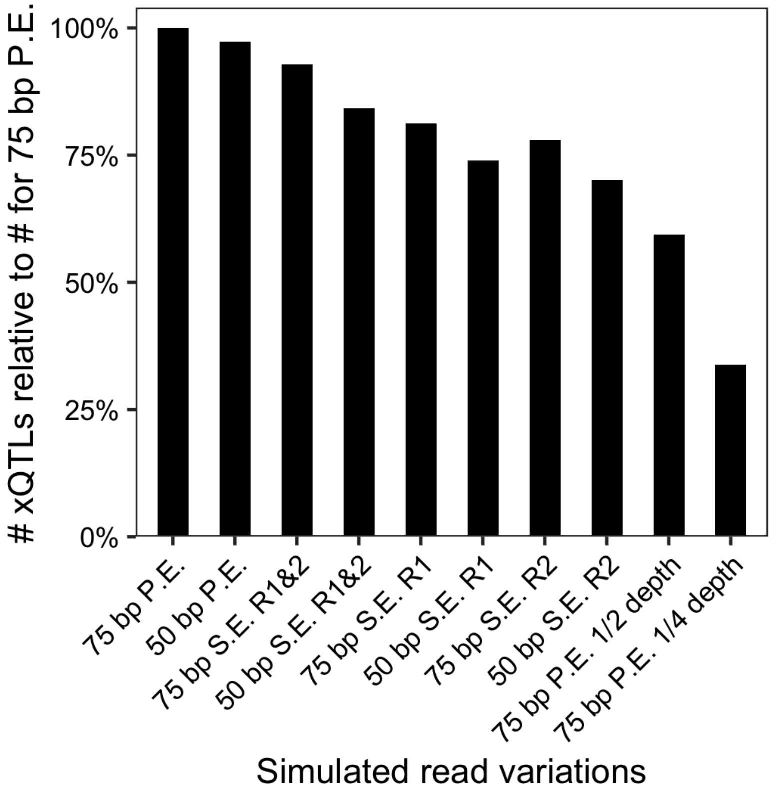


Number of DDP cis-xQTLs when using different variations of the same raw RNA-seq data. A random subsample of 100 Geuvadis samples (75 bp paired-end reads) were truncated to 50 bp, run with R1 and/or R2 reads run as single-end, or run on a subsample of reads equal to a fixed fraction (½ or ¼) of the original reads per sample. The original samples had an average of 30.1 million reads (SD 8.6 million).
